# *De novo* sequencing, diploid assembly, and annotation of the black carpenter ant, *Camponotus pennsylvanicus*, and its symbionts by one person for $1000, using nanopore sequencing

**DOI:** 10.1101/2022.03.31.486652

**Authors:** Christopher Faulk

## Abstract

The black carpenter ant (*Camponotus pennsylvanicus*) is a pest species found widely throughout North America east of the Rocky Mountains. Its frequent infestation in human buildings justifies close genetic examination and its large size and small genome make it ideal for individual sequencing. From a single individual I used long-read nanopore sequencing to assemble a genome of 306 Mb, with an N50 of 565 Kb, and 60X coverage, with quality assessed by a 97.0% BUSCO score, improving upon related ant assemblies. The reads provided secondary information in the form of parasitic and symbiont genomes, as well as epigenetic information. I assembled a complete Wolbachia bacterial isolate with a genome size of 1.2 Mb and 76X coverage, as well as a commensal bacterium of the carpenter ant tribe, the species-specific symbiont *Blochmannia pennsylvanicus*, at 791 Kb, 2400X coverage, which matched to within 200 bp of its previously published reference. I also produced a complete mitochondrial genome with over 5000X coverage, revealing minor rearrangements and the first assembly for this species. DNA methylation and hydroxymethylation was measured simultaneously at whole genome, base-pair resolution level from the same nanopore reads and confirmed extremely low levels seen in the Formicidae family of Hymenoptera. A phased diploid assembly was built, revealing a moderate level of heterozygosity, with 0.16% of bases having biallelic SNPs from the two parental haplotypes. Protein prediction yielded 20,209 putative amino acid sequences and annotation identified 86% matched to previously known proteins. All assemblies were derived from a single Minion flow cell generating 20 Gb of sequence for a cost of $1047 including all consumable reagents. Adding fixed costs for required equipment and batch costs for reagents brings the cost to assemble an ant-sized genome to less than $5000. Complete analysis was performed in under 1 week on a commodity computer desktop with 64 Gb memory.

## Introduction

Complete sequencing and annotation of novel eukaryotic genomes has traditionally only been possible with the support of large institutions and budgets. Previous advancements towards a “$1000 genome” were focused on human genomes that required a pre-existing high-quality assembly and were based on short-read technology. While inexpensive in terms of sample cost, the instrumentation costs remain out of reach of individuals, and require highly trained specialist staff. With the advent of Oxford Nanopore Technologies Minion instrument, DNA and RNA sequencing has become available to individual researchers for accessible costs. Minion flow cells range below $1000 each and have the capacity to sequence up to 30 Gb and the instrument itself retails for under $2000. With minimal additional equipment and reagents, the total cost to build a whole genome sequencing capable lab can be less than $5000. While growing in capability and dropping in cost, the capacity of a Minion flow cell to generate 30X genome coverage, necessary for *de novo* assembly, is currently only feasible for small sized genomes, <500 Mb. Fortunately, one of the most numerous animals on the planet, ants, have both a small genome and are in need of greater genomic resources(1).

The black carpenter ant, *Camponotus pennsylvanicus*, is a prolific pest ranging throughout North America east of the Rocky Mountains and comes into frequent proximity with humans causing significant property damage. Beyond their pest status, the black carpenter ant can reveal biological insights into social behavior, caste determination, and host-endosymbiont interaction, provided we have a high-quality reference genome. For example, within the family Formicidae, a close relative, the Florida Carpenter Ant *C. floridanus*, has been sequenced to high depth using long-read technology(2) and has been used to study plasticity in the biological clock(3), invertebrate calcium regulation(4), and epigenetic control of caste(5). Within the Camponotus tribe, a unique instance of endosymbiosis has occurred where a species of Blochmannia bacteria has become an obligate mutualist and has begun evolving in parallel to the host(6). Additionally, this ant, among many Hymenoptera, hosts endosymbiotic Wolbachia bacteria and can improve the study of this prolific zoonotic(7). Caste determination is still under investigation in Camponotus, which warrants examination of the epigenome. Here we provide base-level measurements of DNA methylation and DNA hydroxymethylation in *C. pennsylvanicus*.

A reference genome for the black carpenter ant will enable study of its ecology, life history traits, and evolution. The ease and low expense of assembly creation and analyses opens avenues to other researchers for this ant and other small-genome insects.

## Results

### Sequencing

I extracted DNA from a single individual worker ant which yielded ~3 μg of total DNA visualized on a gel with high molecular weight, >10kb, but with high fragmentation (Supplementary Figure S1). DNA was sequenced using a single Minion flow cell. Base calling was performed using a GPU optimized version of Oxford Nanopore’s Guppy base caller. Base calling yielded 6.4 million reads and 20.7 Gb of sequence with an N50 of 5113 bp (Table 1.)

**Table 1.**
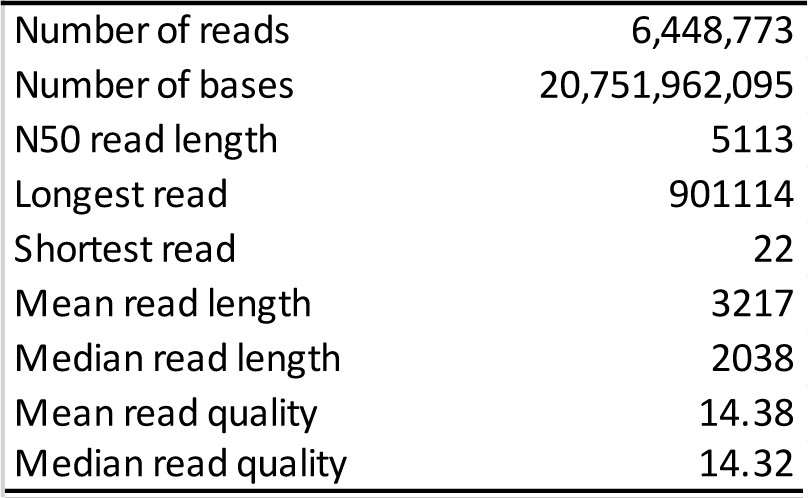
*C. Pennsylvanicus* Read Summary.

### Assembly and filtering

Since no single standard exists for long-read assembly, I performed a comparative analysis on assembly and polishing pipelines and assessed efficacy by BUSCO scoring (Table 2)(8). The genome sequence of a closely related species, the Florida carpenter ant (*Camponotus floridanus*) was used as a benchmark. There are two available assemblies for *C. floridanus*, one created with Illumina short reads (C_flo 1.0) and one with PacBio long-read sequencing (C_flo 7.5). Both assemblies have >50X coverage and greater than 96% BUSCO scores. In order to create an unbiased assembly for *C. pennsylvanicus*, I chose a *de novo* approach, without aligning to any existing data set. I first tried the Shasta assembler since it requires low memory overhead and benchmarks with much faster speed than alternatives. Shasta assembled the genome in less than 1 hour on a single machine. The assembly was sized at 281 Mb with 3,277 contigs, and an N50 of 430 Kb, however the assembly BUSCO score was only 94.1%. Next, I assembled with Flye, which created a 272 Mb assembly and a 96.8% BUSCO score, which was an improvement over the benchmark *C. floridanus* reference genome even before polishing.

**Table 2.**
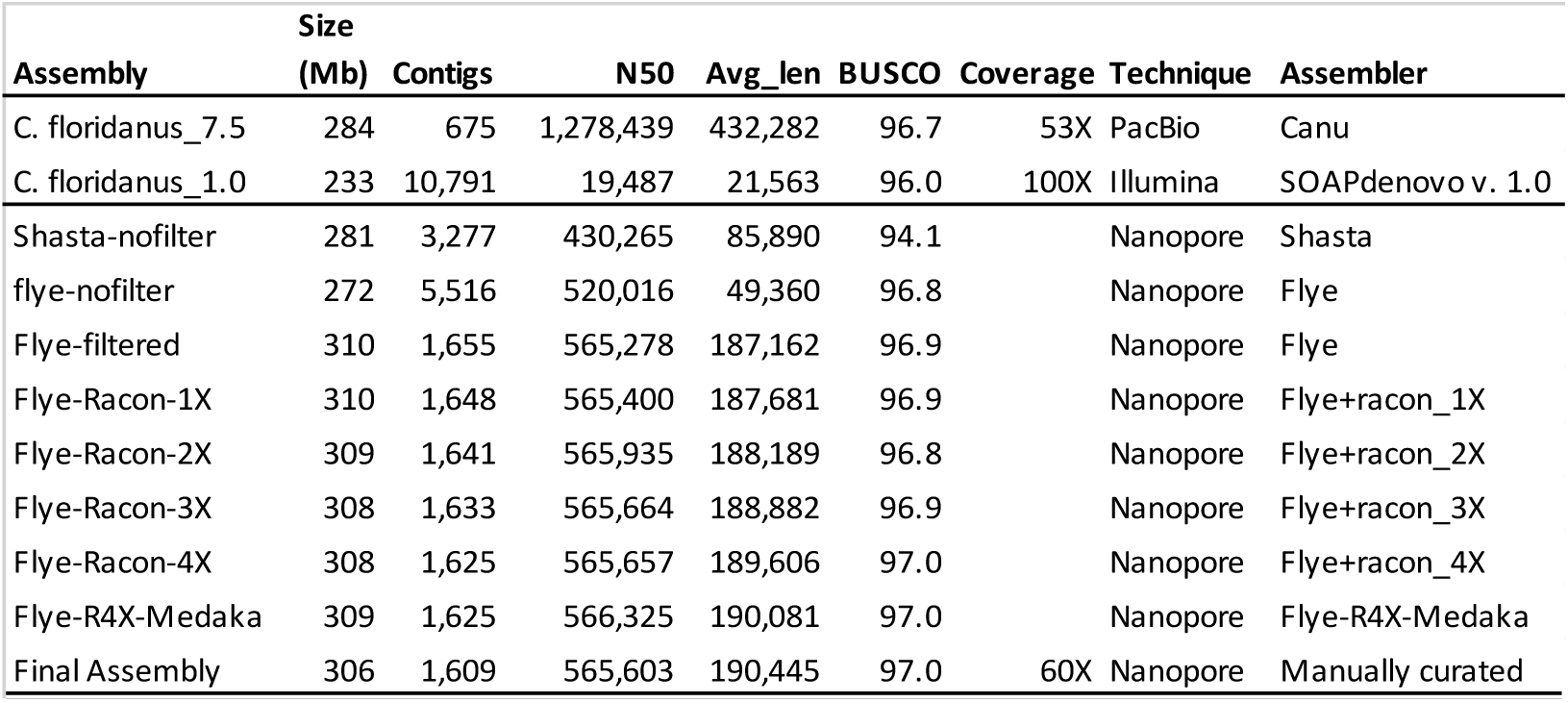
Assembly Statistics.

**Table 2.** Repetitive Element Content of *C. pennsylvanicus*.

|  | Element counts | Total length (bp) | % of Genome Covered |
| --- | --- | --- | --- |
| **Total Bases Masked** |  | 104678206 | 34.16 |
| **Retroelements** | 30521 | 21771960 | 7.11 |
| SINEs: | 2025 | 500013 | 0.16 |
| Penelope | 3155 | 1490951 | 0.49 |
| LINEs: | 10719 | 6928742 | 2.26 |
| CRE/SLACS | 152 | 18220 | 0.01 |
| L2/CR1/Rex | 1114 | 719754 | 0.23 |
| R1/LOA/Jockey | 3564 | 2525973 | 0.82 |
| R2/R4/NeSL | 120 | 35432 | 0.01 |
| RTE/Bov-B | 2232 | 1692132 | 0.55 |
| LTR elements | 17777 | 14343205 | 4.68 |
| BEL/Pao | 2379 | 1846404 | 0.6 |
| Ty1/Copia | 1053 | 376038 | 0.12 |
| Gypsy/DIRS1 | 14030 | 11817732 | 3.86 |
| Retroviral | 46 | 51129 | 0.02 |
| **DNA transposons** | 101469 | 36141816 | 11.79 |
| hobo-Activator | 7468 | 2780257 | 0.91 |
| Tc1-IS630-Pogo | 44001 | 15963966 | 5.21 |
| PiggyBac | 2021 | 228011 | 0.07 |
| Tourist/Harbinger | 2099 | 514354 | 0.17 |
| Mirage | 782 | 389406 | 0.13 |
| Rolling-circles | 5504 | 3278131 | 1.07 |
| Unclassified | 102298 | 33140531 | 10.82 |
| **Total interspersed elements** |  | 91054307 | 29.71 |
| Small RNA | 2098 | 572887 | 0.19 |

Assembly improves dramatically if smaller, lower quality reads are filtered out. For example, Flye with the full unfiltered reads generated a 5,516 contig assembly with an N50 of 520kb. However, when using only half as many reads, by filtering for reads >5kb and dropping the 10% of reads with the worst quality, Flye generated an assembly with only 1,655 contigs, and an N50 of 565kb, despite having nearly identical BUSCO scores. Therefore, I chose to use the filtered read set assembly with Flye for further stages. Subsequent polishing steps used the complete read set.

Four rounds of Racon polishing yielded an assembly with the highest N50 and BUSCO score and was polished a final time with Medaka. After Medaka, the BUSCO score improved to 97.0%. The Flye-Racon4X-Medaka-assembly was 309 Mb in length spread over 1625 contigs. This nanopore-only assembly had the second highest BUSCO score compared to all other currently available ant assemblies, even those produced with hybrid short and long-reads (Figure 1).

**Figure 1.**
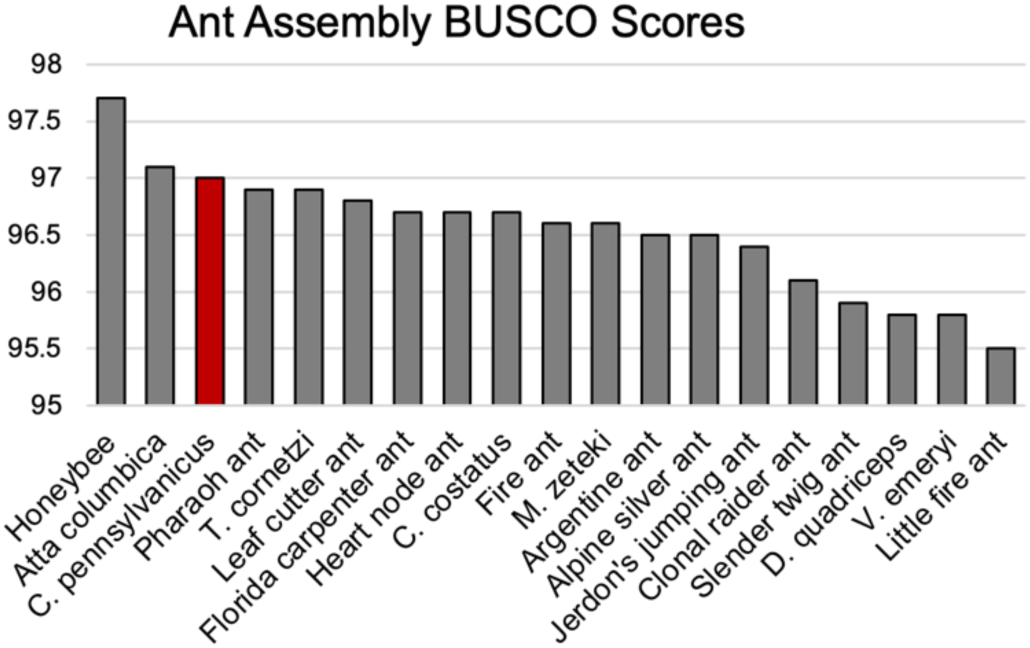
Ant genomes BUSCO scores. *C. Pennsylvanicus* has the second highest ant BUSCO completeness score. BUSCO scores were calculated using the reference genome for at least 1 member of every genus of ant available. Honeybee is included as the most complete hymenopteran available. Two ant assemblies with less than 95% completeness are not shown.

### Contaminants

Careful curation of the assembly is vital to ensure that all contigs represent the true host nuclear genome. I used several measures including GC content, contig coverage, and sequence similarity to known species to filter for any spurious contigs. To identify any known sequence similarity, I first used NCBI’s megablast tool to query the consensus contigs against the NCBI non-redundant nucleotide database (nt). Next, blobtools2 was used to determine the extent of any extra-species inclusion into the assembly contig set (Figure 2). Of the 1625 contigs in the Flye-Racon4X-Medaka assembly, 1580 had a match to an *Arthropoda* species and 31 had no hit to any species and were kept to assemble (Supplementary Table S1). The remaining hits were considered alien contigs with 13 matching to bacterial species and 1 with a small partial hit to an Echinodermata starfish.

**Figure 2.**
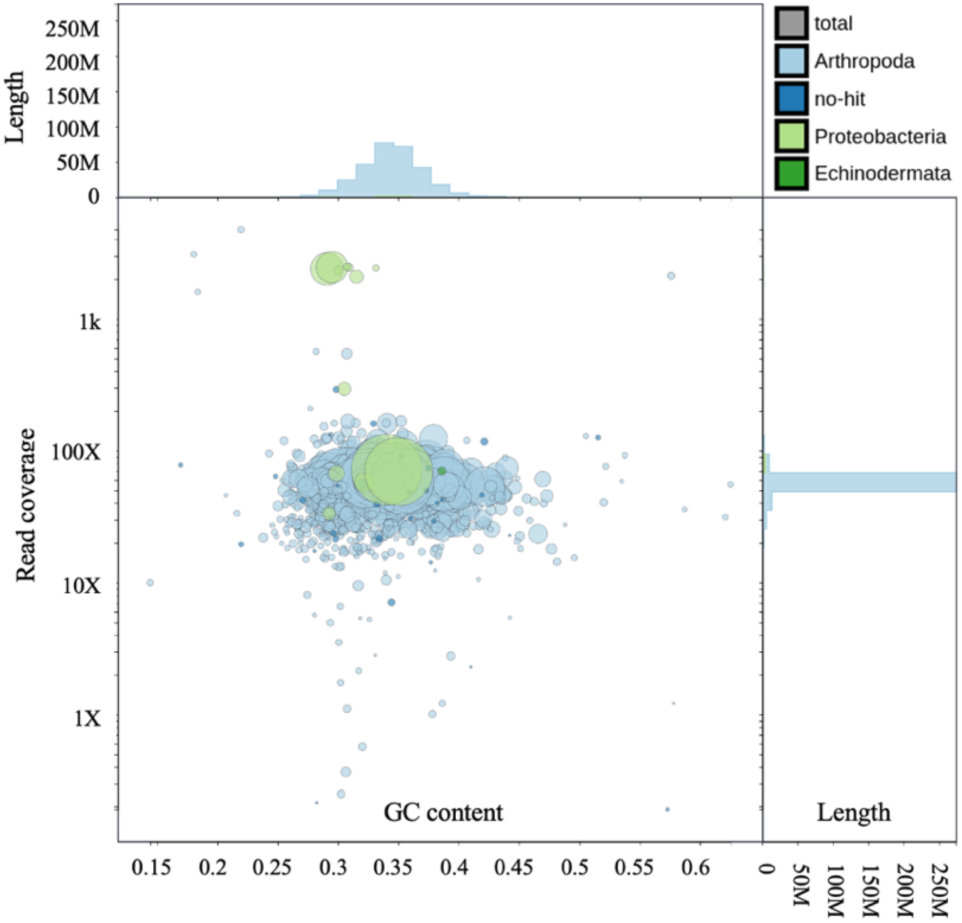
Blobplot of unfiltered consensus contigs. 1625 contigs are shown. Light and dark blue contigs matched to Arthropoda or “no hit” respectively. Higher y-axis blobs have higher coverage. The x-axis is GC content. A single uniform blob indicates single genome origin, rather than multiple species origins. The green blobs with high coverage above the primary blob are of bacterial parasitic origin. The large green blob in the middle is Wolbachia.

To determine origin and filtering criteria of the bacterial and alien contigs, I applied thresholds for identification and removal. Anomalously high coverage proved a reliable marker of non-host contigs. Seven contigs had over 2000X coverage and were of bacterial origin, including nearly the complete genome of *Blochmannia pennsylvanicus*, an endosymbiont of the black carpenter ant. These contigs were removed from the assembly. There were 6 contigs between 30X and 300X coverage that were annotated as bacteria, all *Blochmannia pennsylvanicus*, however close examination of Blast ID showed only fragmentary matches to less than 5% of the query length. The 1 starfish contig was also a less than 5% fragmentary ID match. These were left in the assembly in the assumption that they were misannotated and were true *C. pennsylvanicus* contigs with the following exception. There was a single suspicious contig with typical ant level nuclear DNA coverage. It was a 1.5 Mb contig matching the full length of the *Wolbachia pipientis* genome with no flanking ant sequences. This is consistent with infection rather than an endogenous horizontal insertion, despite this contig having only 69X coverage (Supplementary Figure S2). The list complete list of alien contigs, identification, and removal decision is available in Supplementary Table S2.

The set of 11 contigs with over 1000X coverage that were identified as either “Arthropoda” or “no hit” also proved to be worthy of examination. A group of 3 contigs matched mitochondria genomes from other ant species and were removed. Another was a large rRNA subunit, known to be highly repetitive and thus attract numerous matches collapsed into a single contig. It was kept. There were 5 contigs with anomalously low coverage of less than 1X and they were removed. The manually curated ant assembly contained 1609 contigs.

The generation of so few alien contigs, despite using a whole ant is unsurprising since non-host DNA is unlikely to generate enough consistent coverage to build any consensus fragments. Exceptions were all intracellular high-copy number commensals such as mitochondria, symbiotes and parasites present. However, when assessing contamination, it is important to also examine the complete unassembled read set and not just the assembly. For this purpose, I used the Kraken2 microbiome database. In the ~20 Gb of total reads, a small fraction environmentally derived reads were discovered, with the highest non-ant DNA being human origin at 0.002% frequency.

### Repeat Results

Ant genomes, as with most animals, consist of a large proportion of repetitive sequence made of interspersed transposable elements (TEs) and more simple repeats. Since repeat databases are skewed towards mammals, I chose to use a *de novo* repeat identification pipeline with RepeatModeler to find repeats, and RepeatMasker to classify and annotate them. In total 34.16% of the genome was occupied by repetitive elements, including simple repeats and low complexity regions, at 2.86% and 0.49% of the genome, respectively. Overall, there were 1604 distinct families detected, and not surprisingly, most of the repeats are of DNA transposon origin. Unlike mammals, with retroelement predominant genomes, insect genomes tend to have a greater proportion of DNA transposons(9). All major groups of TEs are distributed in a similar fashion to the TEs described in the Harvester ant (*Pogonomyrmex californicus*) genome(10). To benchmark the *de novo* detected families, I masked the *C. floridanus* genome using the same library and found a slightly lower level, 27.33% of the genome, which may reflect the larger assembly size of *C. pennsylvanicus* (307 Mb) vs. *C. floridanus* (284 Mb), or a more sensitive detection of repeats in my pipeline. The complete list of families is available in Supplementary File S1.

### Diploid genome variation

The genome was approximately 0.16% heterozygous based on biallelic SNPs divided by total callable bases (i.e. the full consensus genome), and 0.21% when based on all heterozygous alleles combined (e.g. indels and SNPs). There were 397,295 transitions (Ts) and 157,601 transversions (Tv) across all the SNPs, leading to a 2.5 Ts/Tv ratio. I generated haploid phased output to better visualize the two haplotypes and their resulting haplotype specific variants such as SNPs and indels. Of 759,295 variants detected by PEPPER-Margin-DeepVariant, most (526,154) were phased, i.e. belonging to a specific parentally derived haplotype. Most variants were also heterozygous. Of 641,913 heterozygous variants, most (482,430) were biallelic SNPs. Examples of phased SNPs and indels can be seen in figure 3. These figures are consistent with a diploid animal genome where most variation between parental haplotypes will be small, e.g. SNVs, followed by larger indels, and fewer inversions, however the variant caller used here does not detect inversions. The use of a single ant allowed parental haplotype resolution, instead of population-based SNP prevalence which would have been produced by pooled samples. A phased variant call file is included with the assembly to distinguish the haplotypes (Supplementary File S2).

**Figure 3.**
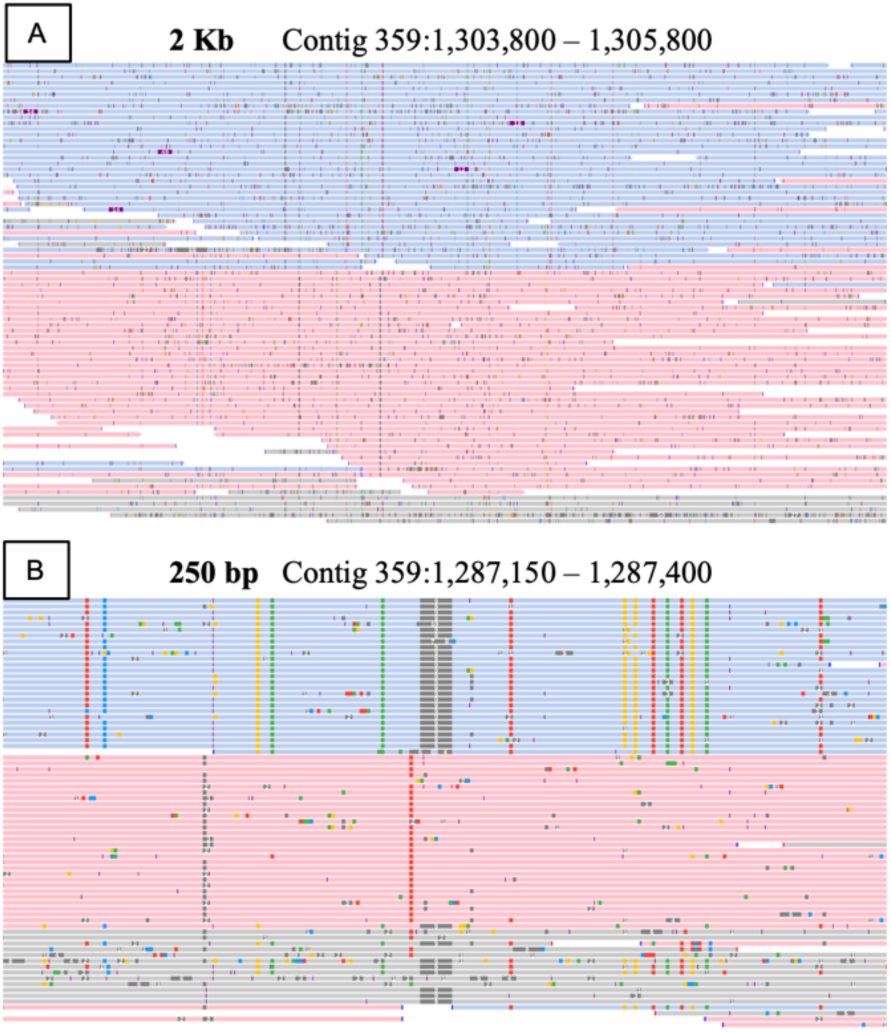
Phased genome view. Depiction of phased reads indicating parental haplotypes at two scales. (A) This 2 kb region illustrates large scale phasing and random sequencing errors. Haplotypes 1 is in blue, haplotype 2 is in red. (B) This 250 bp region illustrates haplotype-specific SNPs and a 9 bp indel in dark gray in haplotype 1. Phased reads show linkage between variants. Reads in gray were not assigned to a specific haplotype.

### Epigenetics

DNA methylation in Hymenoptera is low, generally less than 2% and is not reliably associated with either eusociality or caste determination(11). The nanopore instrument can directly detect DNA methylation and hydroxymethylation at cytosines based on divergent signal intensity compared to unmodified cytosines. Megalodon was used to detect cytosine modifications in all contexts. Here in *C. pennsylvanicus* at CpG cytosines, DNA methylation was 0.359%, and at non-CpG cytosines 5mC was 0.003%. The 5mC level detected by nanopore correlates well with Bewick et al.’s study using whole genome bisulfite sequencing.(11) There are no previous reports describing genome-wide, base-level hydroxymethylation in insects. Here, at CpG cytosines, DNA hydroxymethylation was 0.999% and at non-CpG cytosines 5hmC was 0.101%. With all measures of cytosine methylation and hydroxymethylation below 1%, or near 0%, the detected modified bases are not likely to be biologically relevant.

### Annotation

*Ab initio* protein prediction was performed using Augustus against the masked and non-masked consensus assembly to compare native and transposon derived protein contributions to the transcriptome. There were 20,209 amino acid sequences, representing proteins, detected in the masked genome, made up of 99,891 exons. To determine the identity of these proteins, I compared their sequences to the NCBI ‘nr’ non-redundant database which combines protein sequences from GenPept, Swissprot, PIR, PDF, PDB, and NCBI RefSeq. Of the predicted transcripts from the masked genome, 17,419 identity matches were found, representing 86% of the putative proteins identified by Augustus, suggesting high accuracy and a low false positive rate.

Without masking, there were 40,077 transcripts detected, made up of 150,404 exons. Of the transcripts, 33,286 were identified when compared to the ‘nr’ database, a match rate of 83%. This suggests that the majority of transposon derived transcripts were also positively identified. All the transcripts and IDs are found in Supplementary files S3 & S4.

### Complete C. pennsylvanicus genome deposited

The final assembly of the *C. pennsylvanicus* had a length of 306,426,343 bp, spread over 1609 contigs, with an N50 of 565,603bp, and an average coverage of 60X. A snail plot describing contig length and coverage is shown in Figure 4. The GC content was relatively low for an animal at 34.45%, but in line with other hymenoptera(12). The CG to GC ratio is a nearly even 1.2, reflecting the lack of DNA methylation and concomitant lack of accelerated CpG deamination loss seen in mammals. The diploid assembly was deposited as BioProject PRJNA820489 for the primary haplotype and PRJNA821232 for the secondary haplotype assembly.

**Figure 4.**
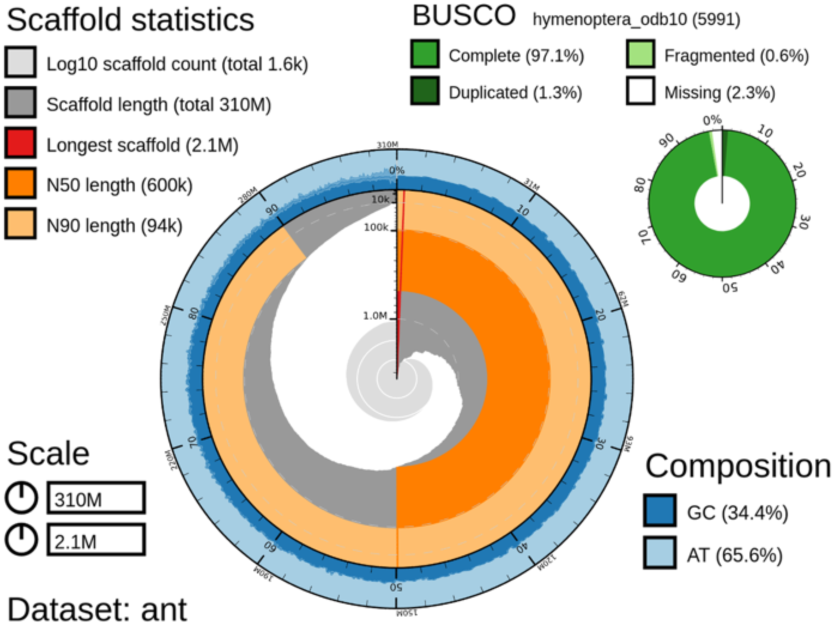
Snail plot of the unfiltered *C. pennsylvanicus* assembly. GC content is shown in the outer ring. The N90 and N50 scores and sizes are shown in light and dark orange respectively.

### Mitochondrial genome

The total read fastq file was also mapped against the mitochondrial genome of the red fire ant (*Solenopsis invicta*, NC_014672.1) to identify reads that were likely mitochondrial in origin. There were 89 Mb of sequence mapping to the reference mitogenome. Resulting hits were extracted, assembled and polished as before. The mitochondrial genome of the black carpenter ant is 16,536 bp, and was covered to 5,385X (Figure 5). It was substantially larger than the 15,549 bp fire ant reference but close in size to the pharaoh ant (*Monomorium pharaonis*) and little fire ant (*Wasmannia auropunctata*) mitogenomes. During annotation 1 frameshift SNP was discovered in a homopolymeric ‘t’ stretch and was manually deleted to create a contiguous coding region. The gene order was broadly similar to mitogenomes in other ants, however several tRNA genes were in different order than even the most similar species available on NCBI’s assembly database. The most similar species were the fire ant and the pharaoh ant. The *C. pennsylvanicus* mitogenome created here was deposited as part of BioProject PRJNA820489.

**Figure 5.**
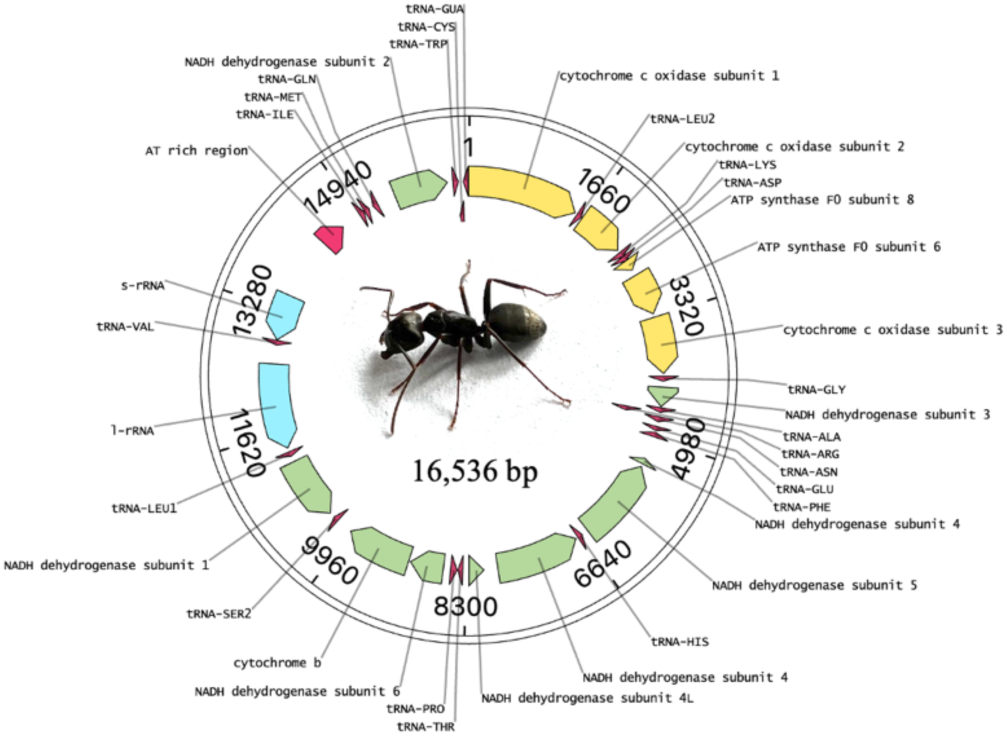
Mitochondrial genome of *C. pennsylvanicus*. The circular length of 16,536 bp was assembled with 5,385X coverage.

### Commensal bacteria B. pennsylvanicus

The tribe of Camponotus ant species have a unique and unusual relationship with the *Blochmannia* genus of endosymbiotic bacteria, without which they do not develop properly. Unsurprisingly then, I found high coverage of reads specifically matching *Blochmannia pennsylvanicus* which has a publicly available sequence. To create a complete assembly of *B. pennsylvanicus* from this ant, I mapped the total reads to the published *B. pennsylvanicus* reference. A total of 192 Mb of sequence mapped to the reference. As with the nuclear genome, Flye was used to assemble a consensus, followed by two rounds of Racon polishing and one round of Medaka polishing. The consensus sequence was 791,499 bp long, with 2,434X coverage. My assembly was within 200 bp of the reference length. It matched the reference genome (NC_007292.1) with 99.84% identity over 100% of the length. It was deposited as BioProject PRJNA821249.

### Wolbachia

Ants, along with the majority of arthropods, are parasitized by Wolbachia bacteria. Within the nuclear assembly, I found and removed a full length Wolbachia genome, and targeted it for separate assembly. A total of 119 Mb of sequence mapped to the *Wolbachia pipientis* reference. Flye, Racon, and Medaka were used to assemble and polish as before. The strain I assembled here had a 1,581,554 bp genome and had only 75X coverage, indicating a 1:1 ratio of Wolbachia to ant cells. My assembly was approximately 400 kb longer than the *W. pipientis* reference. It was deposited as BioProject PRJNA821251.

### Costs and Processing time

As well as expense for the sequencing, time is an important consideration in both sequencing and in data processing. Given the constraints of one person, $1000 in consumables, and one individual ant sequenced, it is appropriate to limit computational hardware to maintain the accessibility of the pipeline described here. Therefore, I restricted all the analyses to using only commodity hardware housed locally in a single desktop computer. The computer had 64 Gb of memory, 4 Tb of SSD storage space, with a 12 core AMD Ryzen 3900x processor and an NVIDIA RTX 2080 Ti GPU to accelerate base-calling and other processing stages.

Costs for sequencing were reduced by using classroom grade equipment that is also suitable for fieldwork. Consumables totaled $1,748 and amortized to $1,047 per sample when accounting for multiple use reagents and kits (Table 3). The largest expense was the flow cell, followed by the library preparation kit. Fixed costs included all equipment necessary for DNA extraction to library preparation, sequencing, and data analysis. The Minion sequencer instrument and the GPU processor were equally expensive at $1000 each. The NVIDIA GPU is required to reduce the base calling stage by several orders of magnitude in time to process. It also accelerates Medaka polishing and variant calling with PEPPER-Margin-DeepVariant. The Guppy base caller requires a minimum of 8 Gb of GPU memory with current neural network models which governs the choice of GPU. A computer running Linux with a minimum of 1 Tb SSD is also required and is assumed to be pre-existing equipment. Optionally the addition of a nanospectrophotometer aids in determining DNA concentration and purity. These instruments can be sourced for less than $5000.

**Table 3.**
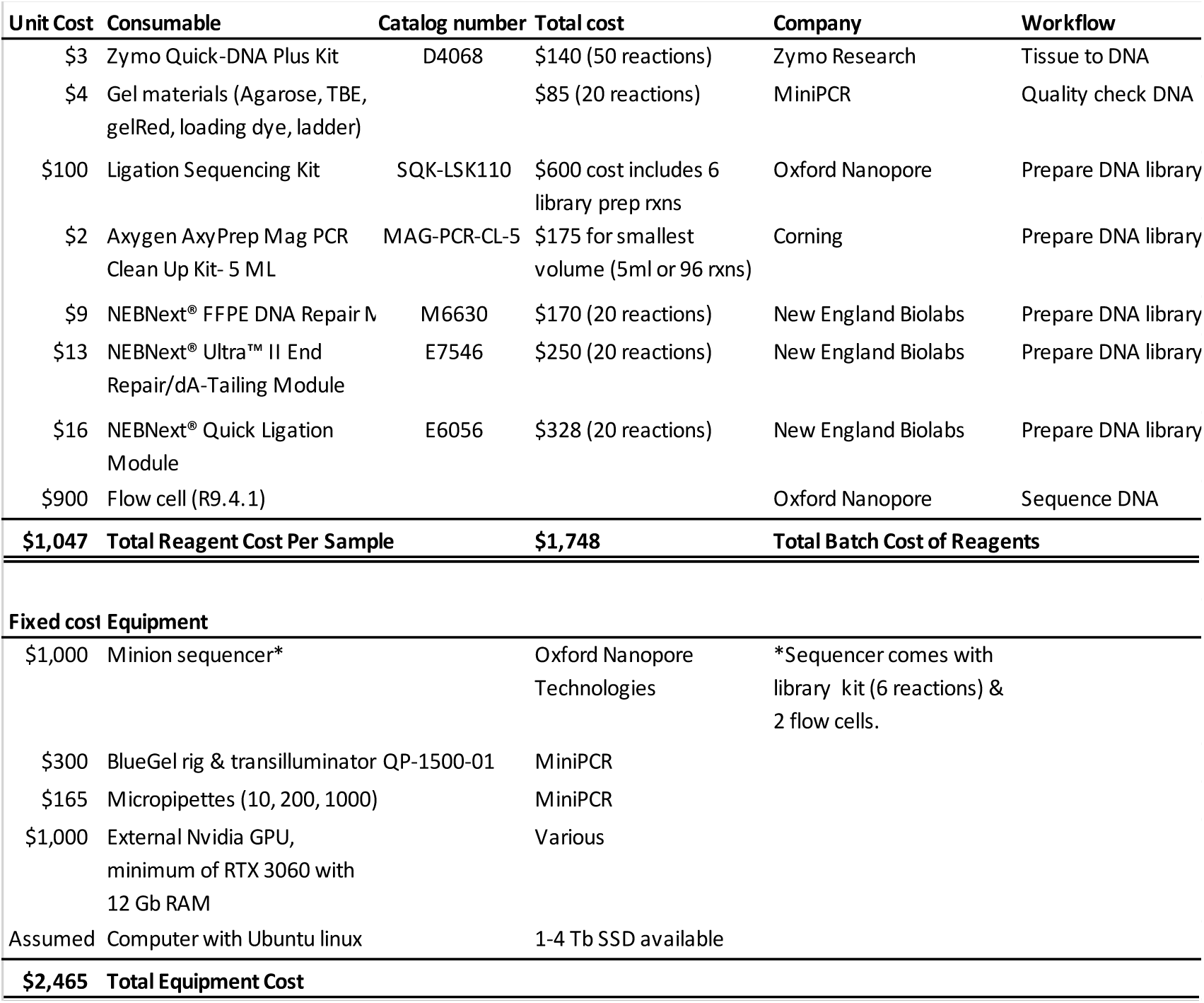
Reagent and Equipment Costs.

The sequencing took 72 hours on an Oxford Nanopore Minion instrument using an r9.4.1 flow cell with a nuclease flush and library reload every 24 hours. Base calling took 6 hours using Guppy software. Assembly with Flye took 2 hours, followed by 4 rounds of Racon and 1 round of Medaka, totaling approximately 6 hours. Repeat identification was the longest stage, taking 28 hours for RepeatModeler2 and 1 hour for RepeatMasker. Modified base calling for DNA methylation took 10 hours with Megalodon. Annotation with Augustus took 6 hours. Variant calling and read phasing took 4 hours with PEPPER-Margin-DeepVariant. The data processing pipeline was performed by the author in less than a week.

## Discussion

### Rationale

Along with the black carpenter ant, several other insect genomes have benefitted greatly from long-read sequencing(13–16). Single insect sequencing has also been previously performed. A single Drosophila fly individual has been sequenced to full genome status was created with great success by using a combination of long and short read sequencing with 3D conformation to bring the consensus to chromosome level(17). Similarly, sequencing 101 Drosophila species using only nanopore reads has also been achieved, revealing many facets of evolution in insects(18). However, these achievements still required large teams and funding. With advancements in nanopore base calling accuracy, cost reductions, and assembly and analysis tools, a $1000 assembly from a single insect and sole researcher has now become possible.

### Assembly

Assembly methods are improving rapidly, as indicated by numerous benchmarking studies(19–21). For small scale labs, a major bottleneck is the computational power needed to assemble a genome from reads. Assemblers working on mammalian size genomes at 3 Gb tend to require high memory, on the order of 2 Tb of RAM. While such machines exist, they are currently out of price range for most labs and available only at high performance computing facilities. Time is also an issue. If using a high accuracy assembler like Canu, even small genomes take two orders of magnitude longer than a nearly equivalent consensus generated with Flye. For instance, a ~10 Mb genome at 70X coverage takes over 22 hours with Canu and less than 1 hour with Flye(22). Mammalian scale assemblies can take weeks to months with Canu. The black carpenter ant genome, at 300 Mb, was able to be assembled in reasonable time frame on consumer level hardware, taking just 2 hours using Flye. Progress towards faster, equally accurate assemblers such as Shasta(23), combined with falling prices of high memory computers will make human scale assemblies available even to small scale labs in the near future.

It is critically important to determine whether the assembly is of high quality before undertaking additional analyses as it will become the publicly available reference genome of this species. Long-reads generated by nanopore sequencing are generally random in errors but do contain some systematic errors that can be overcome through polishing. Vaser et. al recommend 4 rounds of Racon polishing followed by Medaka polishing(24), which is what yielded the best results here. Evaluating assemblies beyond read lengths and quality scores has been greatly enhanced through the benchmarking of universal single copy orthologs (BUSCO) scoring program(8). The BUSCO score is a more biologically relevant determination of whether an assembly is accurate since it detects genes that should be present, rather than relying simply on N50 size or contig number. My comparison reveals that a nanopore-only assembly provided equal or better BUSCO scores than hybrid assemblies of other ant species.

### Contamination

Removal of contamination is vital to generate an accurate public reference genome. Sequencing of whole animals will result in inevitable contamination through a variety of sources. In this case, a wild ant was captured from the environment and may host microbial contamination. Consensus coverage was the most useful filter of non-host DNA. There was a gradation of contig coverage from the mean, 60X, to a high of 566X. Above that range, only 11 contigs exist, and all were above 1600X coverage. Therefore, a cutoff above 1000X was reasonable to identify alien contigs that are not likely to be nuclear in origin. Due to the use of a long-read assembly, contigs should be made of reads derived from single molecules that have substantial overlap. The 5 contigs below 1X coverage imply chimeric joining of unrelated reads into a false contig, so were also removed. GC content has also been used to identify contaminants which often diverge from nuclear genome GC percentage, however in this case, no contigs were flagged as divergent. The use of the Blobtoolkit allowed easy visualization of contig coverage vs. GC content(25).

Finally, during the process of DNA extraction and library preparation, the sample may become contaminated with human or lab-sources of DNA. Cross-checking raw reads using kraken2 efficiently detected human contaminants and detects microbiome reads simultaneously(26).

### Repeat Results

There is a paucity of information on repetitive DNA content in ants. Transposons, retroelements, and simple repeats, are major components of all animal genomes, yet they are not well described in repeat databases. For this reason, it is important to use *de novo* methods of repeat identification before masking the genome to obtain the most accurate percentage of repeat content. Less accurate identification may explain the lower percentage found in the Florida Carpenter ant (27%) vs. its close relative, the black carpenter ant studied here (34%). However, repeat element content can be low in ants, with two lineages of the heart node ant, *Cardiocondyla obscurior*, both having 21% repeat content(27).

### Diploid genome variation

An important advancement will be to develop diploid-aware assemblers to better incorporate heterozygosity seen in diploid species and pooled samples(28). The use of a single ant here allowed detection of phased haplotypes of parental origin based on the biallelic nature of SNPs and indels. The variant call format (vcf) file associated with the primary consensus provides alternative haplotype information, encompassing the full diploid variation of this genome. Phasing reveals linkage of variants from parental haplotigs.

Heterozygosity has traditionally been assessed with only a small number of highly variable markers due to technical constraints, so little data is available to compare full genome wide SNP heterozygosity as I was able to do here. Increasingly measures of whole genome heterozygosity are being adopted. When compared to 101 different drosophila species also sequenced by nanopore and calculated the same way, the SNP heterozygosity of the black carpenter ant (0.16%) fell in the lower middle of Drosophilid range of 0.00035% to 1.1%(18). The SNP transition transversion ratio was found to be 2.5, higher than the 2.1 seen in humans and most mammals(29). This is a natural outcome of mammalian genome methylation at CpG sites which has lead to rapid evolutionary loss of CpG sites, resulting in fewer opportunities for transitions to occur in mammals today. Thus, our ant, without DNA methylation, retains a nearly equal CG to GC ratio of 1.2, unlike humans with 0.25 CG:GC.

### Methylation

Here I replicated the previous reports of extremely low DNA methylation at a base pair specific resolution, and I am the first to report the nearly complete lack of DNA hydroxymethylation in this animal. The lack of DNA methylation was unsurprising, given the generally low levels seen in other studies of Formicidae(30). DNA methylation in the honeybee is known to partially govern caste development, though its level and activity are widely variable in other Hymenoptera, and do not generally correlate with caste development or eusociality(11). For example, in the Florida carpenter ant, histone acetylation appears to be the dominant driver of caste development as seen by Simola et al(5). DNA methyltransferase proteins (DNMTs) do appear to be conserved across Hymenoptera species, however, and DNMTs can have essential functions despite the lack of CpG methylation in other insects(31). An important limitation here is that these results were limited to a single worker, and final determination of the role of DNA methylation in caste development should include queens and other castes in this species.

### Annotation

The Augustus pipeline performed well, identifying the nearly the expected number of 20,000 genes. Some 86% of putative transcripts in the masked genome matched to previously known genes as well as a majority of the transposon ORFs in the unmasked genome. Using only long-read data, without hints from RNA-seq or other species’ expression was a deliberate choice to assess *ab initio* gene modelling in anticipation of this method’s use for less well-studied genera.

### Mitochondrial genome

The full mitogenome was extracted and sequenced marking the first available reference mitogenome for this species. The layout was similar but not exactly convergent with other ant species. Previous nanopore sequencing has recovered mitogenomes from even highly degraded samples such as primate feces, and generated accurate assemblies and so was expected here(32). Copy number of the *C. pennsylvanicus* mitochondria was found to be relatively low compared to the nuclear genome with a ratio of 77:1 mtDNA genomes per haploid nuclear genome. This is significantly lower than in human muscle with varies from 3000-6000 copies per cell or the hundreds to thousands of copies per cell seen in drosophila(33, 34). The low number is likely due to the fact that mtDNA is circular and must be linearized before it can pass through a nanopore for sequencing. Therefore, only fragmented or sheared mitogenomes would be present in the reads, with the majority of circular mitogenomes absent from the data set.

### Commensal bacteria

Ants also contain parasites and symbiotes as well as non-nuclear DNA from mitochondria whose genomes can be assembled from nanopore reads(35). Both Wolbachia and *Blochmannia pennsylvanicus* are known symbiotes of the black carpenter ant and were extracted for full assembly, along with the mitogenome. The Wolbachia assembly was surprisingly much longer than the reference *W. pipientis*, though strains are known to vary widely in co-evolution with their insect host. Copy number was approximately 1:1 based on read coverage. This is lower than is seen in Drosophila, that have a mean of 5.3 copies of Wolbachia per cell(36). In other Hymenoptera, the Wolbachia load can vary from less than 1 to over 10 per cell depending on host factors(37).

The black carpenter ant, and all members of the Camponotus tribe, host specialized endosymbionts of the Blochmannia genus that co-evolve with each species. They are known to have especially slow rates of evolution within species, which aids identification. Here we found the highly abundant coverage of a full-length *B. pennsylvanicus* genome in its own contig. It differed by only 200 bp in length from the reference genome with a nearly perfect 99.84% sequence identity, confirming both the species of bacteria as well as the host ant.

### Project cost and processing time

To complete this assembly for approximately $1000, some DNA quality control steps were simplified. Sizing and fragment validation were performed on an agarose gel rather than the recommended Agilent TapeStation or Bioanalyzer(38). The gel rig and pipettes were robust classroom grade equipment available from Amplyus that are less expensive than laboratory grade hardware. Quantitation was performed on a nanophotometer that requires no reagents instead of a more sensitive qubit fluorometer (ThermoFisher Inc.) and still yielded reliable results. Oxford Nanopore Technologies makes the Minion nanopore sequencer and associated flow cells and reagents available to purchase affordable by individual labs. I chose to use the Ligation Sequencing kits SQK-110 for higher throughput, but Rapid Sequencing kits and Field Sequencing Kits can eliminate additional reagents or allow cold-chain free sequencing respectively. Limiting the project to the use of commodity computer hardware makes this pipeline more accessible to small scale labs. The use of classroom grade gel equipment makes sequencing more accessible in the field. Combined, these optimizations put full genome assembly, at least for small size genomes, in reach of a larger share of the scientific community.

## Materials and Methods

### Tissue collection

The author captured a single individual carpenter ant worker (*Camponotus pennsylvanicus*) in the attic structure of a residence in St. Paul, MN on February 18^th^, 2022. Species identification was confirmed by the UMN Extension service. Holotype ant photo is available as Supplementary Figure S3.

### DNA extraction

DNA was extracted using a Zymo DNA plus miniprep kit. The entire worker ant was homogenized using a plastic pestle inside a 1.5 ml Eppendorf tube with 300 μl of Zymo DNA shield. The standard miniprep protocol yielded approximately 3 μg of total DNA. Quality was assessed using a nano-spectrophotometer (Implen N60, Munich Germany) and run out on a MiniPCR bluegel student electrophoresis rig by Amplyus (Cambridge, MA). Size was >10,000 bp with visible fragmentation smearing (Supplementary Figure S1).

### Library Preparation

Library prep was performed with an Oxford Nanopore SQK-LSK-110 kit according to manufacturer’s instructions with the following changes. The total amount of 3 μg of DNA was used in a single library prep reaction and eluted with 45 μl of elution buffer, then split into 3 aliquots of 15 μl libraries to facilitate reloading of the flow cell.

### DNA sequencing

Sequencing was performed using Oxford Nanopore MINKNOW software (v21.11.9) and Guppy (v5.1.15) set to fast basecalling. Post-hoc basecalling was performed using Guppy with the ‘super accuracy’ model (dna_r9.4.1_450bps_sup.cfg).

### Computational Methods

Detailed instructions for the pipeline used here are available as Supplementary File S5. Below are abbreviated methods.

### Genome Assembly

Genomes were assembled with Shasta(23) and Flye(39), followed by rounds of Racon and Medaka polishing. Genome assembly with Shasta v0.8.0 was performed with the following parameters, ‘sudo ~/shasta-Linux-0.8.0 --input ant-total.fastq.gz --config Nanopore-Oct2021 -- memoryBacking disk --memoryMode filesystem --Reads.minReadLength 1000 -- assemblyDirectory ShastaRun’. Genome assembly with Flye v2.9 was performed with the following parameters, ‘flye --nano-hq ant-total.fastq --out-dir flye-results --genome-size 281m -- threads 23’. The size estimate of 281 Mb came from estimates for Tsutui et al. for members of family Formicidae(40). BUSCO v5.2.2 scores were calculated using the following command, ‘busco -i assembly.fasta -o busco -m genome --lineage hymenoptera-c 23’.

### Polishing

Racon was run with the following parameters. ‘racon --cudapoa-batches 40 -t 23 ant-total.fastq.gz ant-total.sorted.sam assembly.fasta > assembly-racon1.fasta’. This step was repeated 4 times. This version of Racon was compiled with CUDA support which sped up processing to under 1 hour per cycle. After each Racon polish step, the raw reads must be re-aligned to the resulting assembly with minimap2 prior to the next polishing step(41). Medaka similarly has a GPU mode which greatly decreases polishing time to less than 4 hours. It was run with the following parameters. Environment variable was set with ‘export TF_FORCE_GPU_ALLOW_GROWTH=true’, and polishing performed with ‘medaka_consensus -b 100 -i ant-total.fastq -d consensus-racon.fasta -o medaka-results/ -t 23 -m r941_min_sup_g507’.

### Contamination removal

NCBI’s megablast was used to determine contig identity. Blobtools2 was used to visualize a density plot of GC content vs. genomic coverage with interactive selection of contigs by phylum and species(25). Blobtools2 local installation was used to generate the blobplot, snail plot, cumulative count, and BUSCO plot as well as filtered table output. The blobtoolkit requires the consensus, coverage statistics generated with samtools, and the blast output with species and taxonomy. Details on configuration are provided in supplementary methods. Alien contigs flagged for removal were deleted manually with the nano text editor.

### Commensal and organelle assembly

Total reads were individually aligned to reference genomes for *Blochmannia pennsylvanicus* (GCF_000011745.1), *Wolbachia pipientis* (GCF_014107475.1), and the mitogenome of the red fire ant (*Solenopsis invicta*, NC_014672.1) using minimap2. Flye required the ‘--meta’ tag for the mitochondrial assembly and the ‘--asm-coverage 50’ flag for the *B. pennsylvanicus* assembly.

### Coverage assessment

Depth statistics were generated with mosdepth v0.3.3(42).

### Repeat identification

Since annotated repeats in insect genomes are sparse, repeat identification requires two stages. First *de novo* repeat identification was performed with RepeatModeler2 v2.0.2a(43). Second, the libraries generated with RepeatModeler2 were used as input to RepeatMasker v4.1.0 to create a complete genome annotation of repeats and classified using existing names for classes, families, and subfamilies from the Dfam v3.5 open source repeat library where known and novel IDs for previously unknown families(43, 44). The family consensus sequences are found in Supplementary File S1, ‘C_pennsylvanicus-repeat-families.fa’.

### Gene annotation

Augustus v3.4.0 was used for *ab initio* protein prediction using the honeybee as nearest species(45). The masked consensus was used for prediction to eliminate false positives from open reading frames present in transposons. To identify the proteins, I used DIAMOND(46) to match predicted proteins against the NCBI non-redundant protein database, ‘nr’. The complete annotation matches are in Supplementary File S4.

### Variants and Diploid Construction

Sequence variants were called against the haploid consensus using the PEPPER-Margin-DeepVariant v0.7 pipeline with GPU acceleration and phased haplotype output(47). Whatshap v1.3 was used to calculate statistics from the resulting ‘vcf’ file(48). The original consensus assembly is considered the primary haplotype. The secondary haplotype consensus was built by swapping biallelic variants from the vcf file into the primary consensus using ‘bcftools consensus PEPPER_MARGIN_DEEPVARIANT_FINAL_OUTPUT.phased.vcf.gz > haplotype2.consensus.fasta’.

### Epigenetic Marks

DNA methylation (5mC) and hydroxymethylation (5hmC) were determined using megalodon v2.4.2 with a base calling model capable of detecting both marks on cytosines in any context natively, res_dna_r941_min_modbases_5mC_5hmC_v001, using the same reads as used to generate the assembly. Post processing yielded files containing 5mC or 5mC marks at CpG sites and CH sites within the consensus assembly.

## Acknowledgements

Thanks to AK Barks for critically reading the manuscript.

## Author contributions

CF captured the ant. CF performed sequencing and all analyses. CF wrote the manuscript.

## Data Availability

DNA reads have been deposited in fastq format in the Sequence Read Archive (SRA) repository. (https://www.ncbi.nlm.nih.gov/sra) under SRA submission SRR18530582. Assembly is available as BioProject PRJNA820489 for the primary haplotype and PRJNA821232 for the secondary haplotype. The mitogenome was deposited as part of BioProject PRJNA820489. B. pennsylvanicus assembly was deposited as BioProject PRJNA821249. Wolbachia species associated with C. pennsylvanicus was deposited as BioProject PRJNA821251

## Supplementary Materials

**Supplementary Figure S1**. Gel image of input DNA.

**Supplementary Figure S2**. Wolbachia blast comparison.

**Supplementary Figure S3**. Holotype ant.

**Supplementary Table S1**. Complete contig list of coverage, id match, GC content, and length.

**Supplementary Table S2**. Alien contig list

**Supplementary File S1**. Repeat families identified by RepeatModeler2.

**Supplementary File S2**. Phased variant call format file.

**Supplementary File S3**. Transcripts in amino acid fasta format.

**Supplementary File S4**. Blast id match to transcript names.

**Supplementary File S5**. Complete computational methods.

**Supplementary Figure S1.**
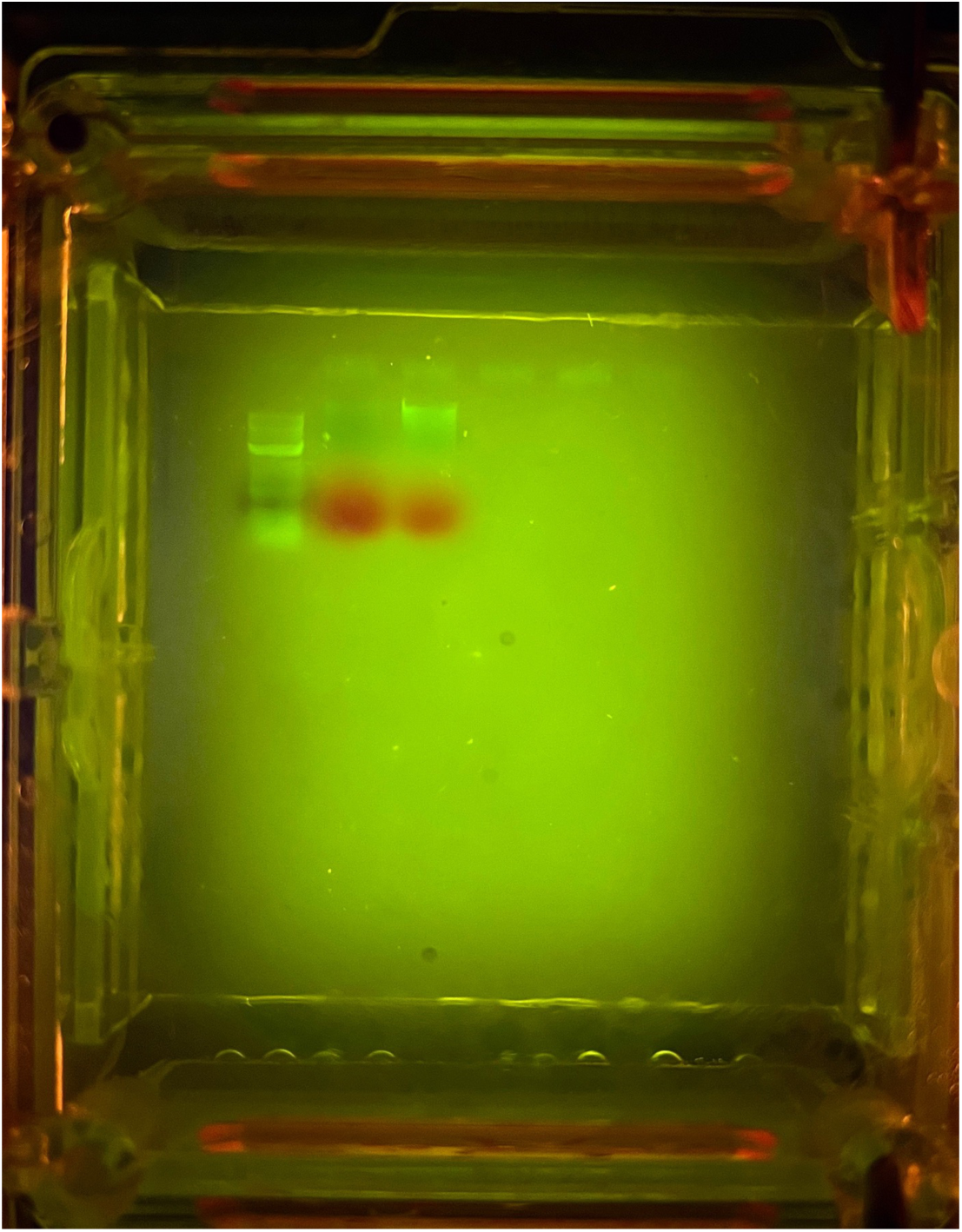
Gel image of DNA DNA Gel image of input DNA. The DNA of a single ant is shown (lane 2). 10 kb ladder indicates high molecular weight but with high amount fragmentation visible as a smear. Visualization was performed on a MiniPCR bluegel student gel electrophoresis rig using safe stain.

**Supplementary Figure S2.**
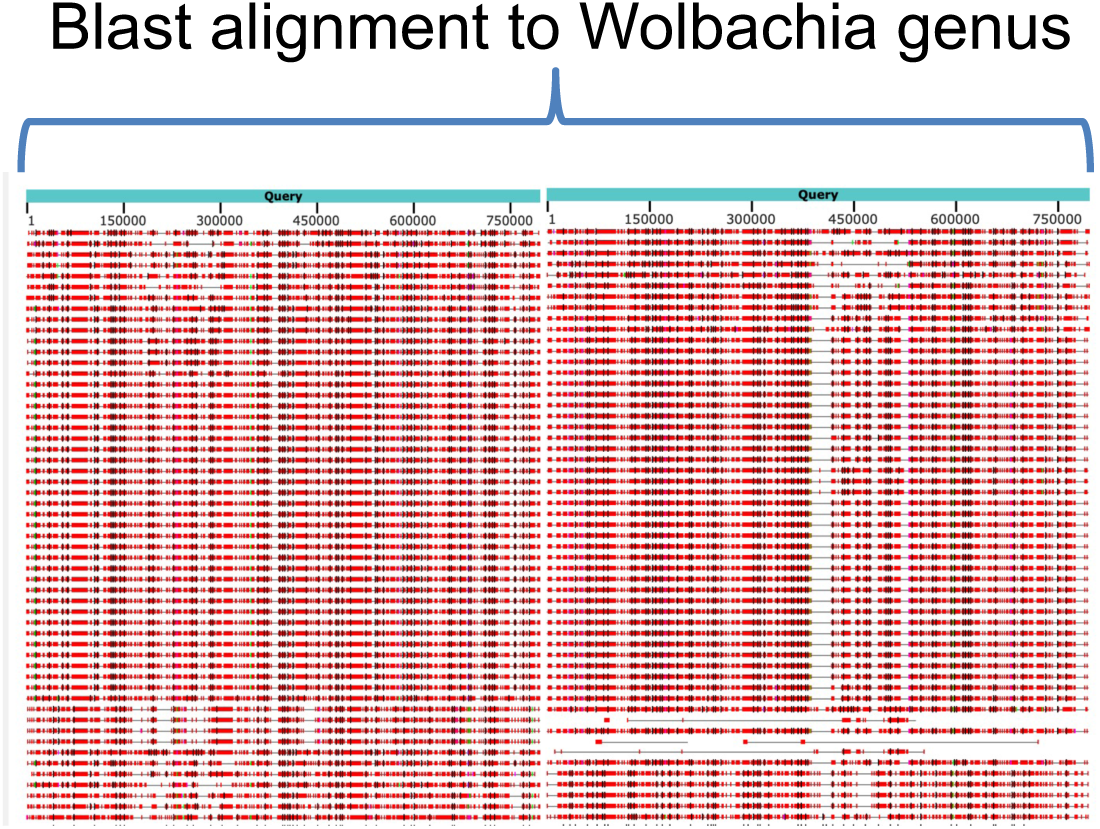
Wolbachia blast Contig_2049 was split into two chunks to fit within NCBI Blast webpage limits for visualization. The contig has hits across the complete length, with no flanking matches outside of Wolbachia genus indicating this is an independent genome and not an instance of horizontal transfer.

**Supplementary Figure S3.**
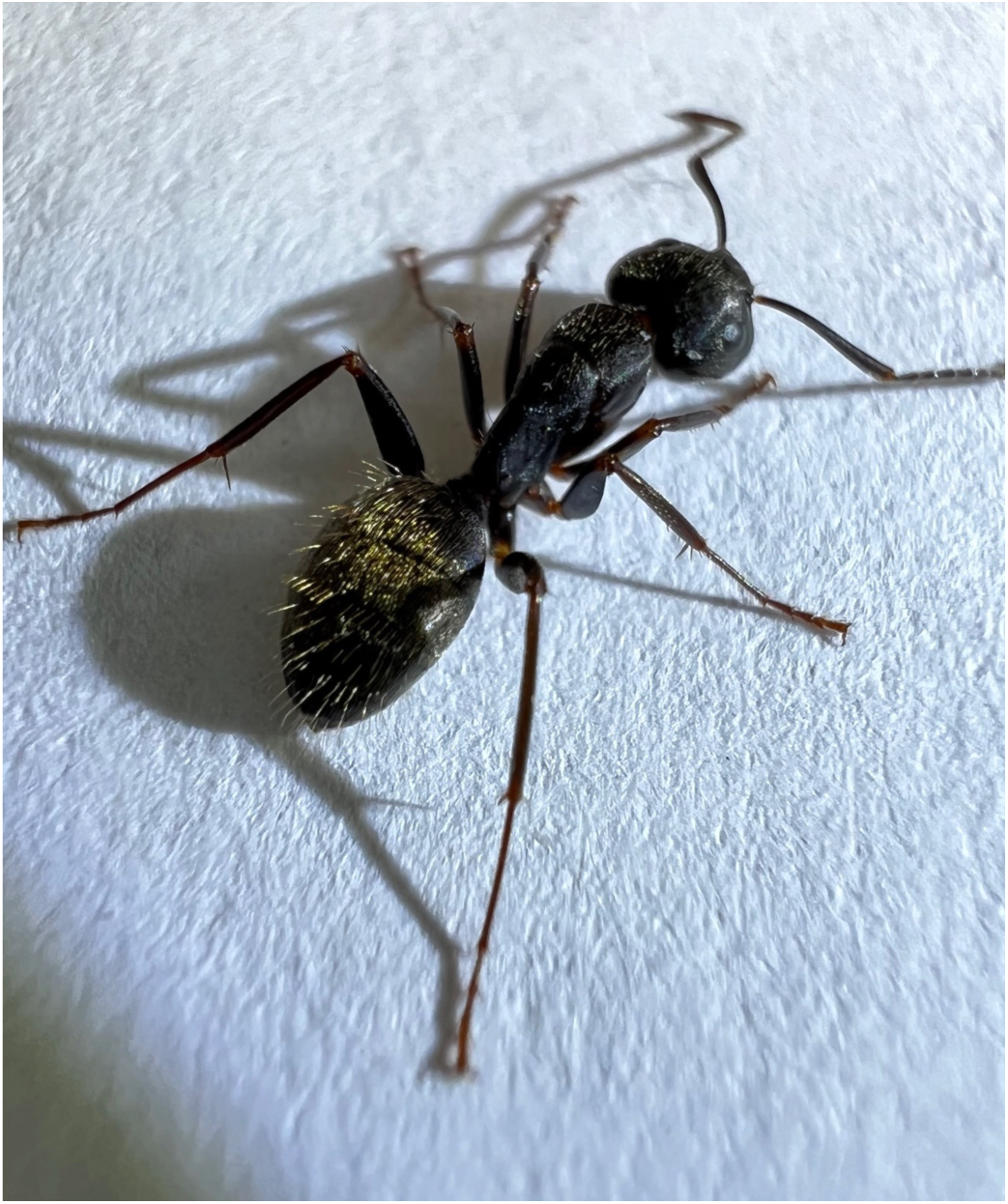
Holotype Ant This is the holotype individual of *C. pennsylvanicus* used for this genome assembly.

