## Supplementary File S5 for "*De novo* sequencing, diploid assembly, and annotation of the black carpenter ant, *Camponotus pennsylvanicus*, and its symbionts by one person for $1000, using nanopore sequencing": Supplementary File S5. Ant-pipeline.nb.html

Camponotus pennsylvanicus Nanopore Sequencing


Code 

- Show All Code
- Hide All Code
- Download Rmd

### Camponotus pennsylvanicus Nanopore Sequencing

###### Chris Faulk

#### 3/2/21


### File S10. Complete Computational Methods

Christopher Faulk (UMN)

### Background

#### The Ant

A diploid reference assembly was built using a single individual
worker of *Camponotus pennsylvanicus* along with mitogenome and
its symbionts, Wolbachia spp. and *Blochmannia
pennsylvanicus*.

#### The Experiment

Sample origin: St. Paul Attic  
Extraction method: Zymo DNA miniprep plus kit.  
Sequencing instrument: Oxford nanopore Minion, flowcell r9.4.1  
Sequencing took place over 4 days with 3 libraries loaded, each preceded
by one wash (wash kit 004).  
Library prep: 3 aliquots of 1 ug of DNA from a single ant sample were
each end prepped using NEBNext FFPE Repair Mix (M6630), NEBNext Ultra II
End repair/dA-tailing Module (E7546), and NEBNext Quick Ligation Module
(E6056) and subsequently library prepped using the ONT supplied SQK-
LSK110 kit.

### Methods

Computational environment and software: Sequencing computer was built
with an AMD Ryzen 3900x processor with 12 cores, 24 threads, 64 Gb RAM
and a 4 Tb SSD, running Ubuntu Linux 20.04. For base calling I added two
GPUs, a GeForce 2080Ti. I monitored the GPU status with
`nvtop`.

#### Run length

The flow cell generated 8.34 million reads over 72 hours.

#### Read generation

Fast5 files were generated from Oxford Nanopore Minion instrument and
called with Guppy v5.1.15. I used the GPU enabled version of guppy in
concert to enable live base calling with the fast base calling model and
for post-hoc base calling I used the super (sup) accuracy model with the
following post-hoc parameters.

`guppy_basecaller --config dna_r9.4.1_450bps_sup.cfg --compress_fastq --device cuda:0 --nested_output_folder --recursive --input_path no_sample/ --save_path ./sup`

### Analysis

The following pipeline was used for analysis.

#### QC with Nanoq

Nanoq gives fastq read statistics from fastq.gz files and has many
other great features and is fast.

**Ant total** ant reads combined:
`nanoq -v -s -i ant-total.fastq.gz`

#### Nanoq Read Summary

**“ant-total.fq.gz**”  
Number of reads: 6,448,773  
Number of bases: 20,751,962,095  
N50 read length: 5113  
Longest read: 901114  
Shortest read: 22  
Mean read length: 3217  
Median read length: 2038  
Mean read quality: 14.38  
Median read quality: 14.32

### Microbial and vertebrate contamination

To determine whether there is contamination in the raw reads, we must
scan the raw fastq file. Kraken2 uses a 16 Gb
database for microbial sequence identification, as well as mouse and
human.

Run kraken2 to identify.  
`kraken2 --db /mnt/bc91d872-d479-41b6-b994-c521c27d41cf/kraken2/k2_pluspf16gb/ --threads 10 --use-names --report ant-total.FASTQ.unmapped.report.txt --output ant-total.FASTQ.unmapped.out.txt ant-total.fastq`

Pavian was used
to visualize. From inside R run
`pavian::runApp(port=5000)`

### Genome assembly

#### Shasta assembly

Genome assembly was performed with Shasta assembler
v0.8.0 due to its faster speed in comparison to Canu and flye. The
following parameters were given, and the input was the total reads fastq
file.  
`sudo /home/cfaulk/Desktop/shasta-Linux-0.8.0 --input ../ant1/sup/ant-total.fastq.gz --config Nanopore-Oct2021 --memoryBacking disk --memoryMode filesystem --Reads.minReadLength 1000 --assemblyDirectory ShastaRunOct2021`

Afterwards run `shasta --command cleanupBinaryData` to
clean up binary data.

#### Flye assembly

##### Filter reads

Assembly contigs improve dramatically if smaller, lower quality reads
are filtered out. For example, using flye on the complete data set
generated a 5,516 contig assembly with an N50 of 520kb. However, when
using only half the reads, filtered for reads >5kb and dropping the
10% of reads with the worst quality, flye generated an assembly with
1645 contigs, and an N50 of 599kb, despite having nearly identical BUSCO
scores. Therefore, I chose to use the filtered read set assembly with
flye for further stages. Subsequent polishing steps used the full read
set.

Filtering was performed with `filtlong` with
the following parameters.  
`filtlong --min_length 5000 --keep_percent 90 input.fastq.gz | gzip > output.fastq.gz`

##### Assemble

Flye v2.9.1 was
used.
`flye --nano-hq ant-total-filtlong-5000-90perc.fastq --out-dir flye-results --genome-size 281m --threads 23`

##### Polishing (Racon + Medaka)

- Racon was run with the
  following parameters. Note that this version of Racon was compiled with
  CUDA support. Remember to re-align the reads to the new assembly after
  each Racon run before the next cycle.
  `racon --cudapoa-batches 40 -t 23 ../ant-total.fq.gz ant-total.racon0.sam flye-results/assembly.fasta > assembly-racon1.fasta`

Realign reads for next round.  
`minimap2 -ax map-ont assembly.fasta ../ant-total.fastq -t 23 > ant-total.racon0.sam`

- Medaka was run
  after 4 rounds of racon (because RaconX4 had the highest BUSCO) It has a
  GPU mode. `export TF_FORCE_GPU_ALLOW_GROWTH=true`  
  `medaka_consensus -b 100 -i ../ant-total.fastq -d racon-results/Assembly-racon2.fasta -o medaka-results/ -t 23 -m r941_min_sup_g507`

#### Identify contigs by NCBI Blast.

First download the nt database from NCBI.

The following blast query will provide correct input format for
blobtools.  
`blastn -query consensus.fasta -task megablast -db ~/Desktop/genomes/nt/nt -outfmt '6 qseqid staxids bitscore std sscinames sskingdoms stitle' -culling_limit 10 -num_threads 23 -evalue 1e-3 -out consensus.fasta.vs.nt.cul5.1e3.megablast.out`

#### Examine assembly for contaminants with Blobtoolkit

Filter the assembly for bacterial reads using BlobTools2.

1. Generate a coverage file:
   `samtools coverage ../racon-medaka-out/consensus-medaka.sorted.bam > ant-total.medaka.sorted.txt`
2. Create the blobdir with the consensus fasta and add the BLAST
   data to it:
   `blobtools create –fasta ../racon-medaka-out/consensus.fasta –meta ant.yaml –taxid 104422 –taxdump taxdump/ ant blobtools add –hits consensus.medaka.fasta.vs.nt.cul5.1e3.megablast.out –taxdump taxdump/ ant`
3. Add the coverage file to the blobdir:
   `blobtools add --text ant-total.medaka.sorted.txt --text-header --text-cols '#rname=identifier,meandepth=ant_reads_cov' ant`
4. Fix the axes:
   `blobtools add --key plot.y=ant_reads_cov ant`
5. Add the BUSCO scores:
   `blobtools add --busco summary.tsv ant`
6. View blobplots in a browser:  
   `blobtools view --local --interactive ant` http://localhost:8007/view/ant/dataset/ant/blob

##### Manually remove contaminants

Look at the csv made by blobtools and figure out which contigs are
alien. Examine and remove contigs greater than 1000X and less than 1X.
Remove them using nano or other text editor and save as a fasta.

#### Consensus read alignment

Now that we have a cleaned, polished consensus assembly, we need to
re-align the reads to it to get final coverage data.
`minimap2 -ax map-ont -t 24 consensus.clean.fasta ../ant-total.fastq.gz > ant-total.mapped.to.consensus.clean.sam`

Convert from sam to bam and sort alignments all in 1 step.
`for i in *.sam; do samtools view -u $i |samtools sort -@ 24 -o $i.sorted.bam ; done`
Index the bam file
`for i in *.bam ; do samtools index $i -@ 24; done`

### Assembly stats

#### Mosdepth

Generate depth statistics overall
`mosdepth -n --fast-mode -t 4 <prefix> file.bam`  
Generage depth statistics for 500 bp windows
`mosdepth -n --fast-mode --by 500 -t 4 <prefix> file.bam`

#### BUSCO

After each assembly or polishing step BUSCO (v5.2.2) score was calculated
in genome mode with hmmsearch v3.1 and metaeuk v5.34. The assembly was
tested for completeness against the hymenoptera\_odb10 database.
`busco -i Assembly.fasta -o busco -m genome --auto-lineage-euk -c 23`
`busco -i Assembly.fasta -o busco -m genome --lineage hymenoptera -c 23`

### Repeat Identification

Since annotated repeats in insect genomes are sparse, repeat
identification requires two stages. First *de novo* repeat
identification was performed with RepeatModler2.
Second, the libraries generated with RepeatModeler2 were used as input
to RepeatMasker to create a
complete genome annotation of repeats with existing names for classes,
families, and subfamilies where known and novel IDs for previously
unknown families.

RepeatModeler was run to identify repeat content. Takes about 24
hours on a 300 Mb insect genome.

1. First build database from consensus
   fasta.`/usr/local/RepeatModeler-2.0.2a/BuildDatabase -name "ant-genome" ../clean-assembly/consensus.clean.fasta`
2. Run RepeatModeler2.  
   `nohup <RepeatModelerPath>/RepeatModeler -database ant-genome -pa 14 -LTRStruct >& run.out &`
3. RepeatMasker was then run to determine genomic repeat content
   specifying the custom library created with RepeatModeler.
   `RepeatMasker -pa 20 -s -xsmall -lib ant-genome-families.fa ../clean-assembly/consensus.clean.fasta`

### Genome Annotation

Augustus 3.4.0 was used for *ab initio* protein model
annotation.
`augustus –species=honeybee1 ../clean-assembly/consensus.clean.fasta.masked –softmasking=1 > augustus-predictions-softmasked.gff –progress=TRUE`

Next, extract the nucleotide and amino acid hits from the gff file
into fasta files.
.`/augustus-master/scripts/getAnnoFasta.pl augustus-predictions-softmasked.gff –seqfile=../clean-assembly/consensus.clean.fasta >20209 amino acids detected in augustus-predictions-softmasked.aa`

#### Identification of Augustus protein models

Get the NIH nr (non-redundant protein database for blastp
searching)

1. `wget 'ftp://ftp.ncbi.nlm.nih.gov/blast/db/nr.*.tar.gz'`
2. `cat nr.*.tar.gz | tar -zxvi -f - -C .`

Use DIAMOND to speed up matching over blastp which takes forever.

1. `diamond -prepdb -d nr`
2. `diamond blastp --query augustus-predictions-softmasked.aa -d ~/Desktop/genomes/nr/nr -p 22 -v -o augustus-predictions-softmasked.diamond.out --max-target-seqs 1 -f 6 qseqid sseqid pident length mismatch gapopen qstart qend sstart send evalue bitscore slen nident mismatch stitle salltitles`

### Diploid assembly

A diploid assembly provides calls for both parental haplotypes,
yielding essentially two assemblies with phased haplotype blocks. This
is the future of assemblies rather than the traditional haploid
assemblies that collapse heterozygosity into a single “canonical” base.
However these require ~60x coverage, which we have with a single ant
sample.

#### Call variants

We used an updated version of polishing and variant calling from the
manuscript Shafin et al. “Haplotype-aware variant calling with
PEPPER-Margin-DeepVariant enables high accuracy in nanopore long-reads.”
Requires docker and nvidia-docker toolkit. Takes about 1 hour.

**Command specifying directories with Docker.** Finds
variants, e.g. generates a single consensus + vcf file.
`sudo docker run -it --ipc=host --gpus 1 -v "/home/cfaulk/Desktop/ont-sequencing_runs/ant-analysis/pepper-margin-deepvariant-out/:/home/cfaulk/Desktop/ont-sequencing_runs/ant-analysis/pepper-margin-deepvariant-out/" kishwars/pepper_deepvariant:r0.7-gpu run_pepper_margin_deepvariant call_variant -b /home/cfaulk/Desktop/ont-sequencing_runs/ant-analysis/pepper-margin-deepvariant-out/ant-total.mapped.to.consensus.clean.sorted.bam -f /home/cfaulk/Desktop/ont-sequencing_runs/ant-analysis/pepper-margin-deepvariant-out/consensus.clean.fasta -o /home/cfaulk/Desktop/ont-sequencing_runs/ant-analysis/pepper-margin-deepvariant-out/pepper-margin-deepvariant-out -t 18 -g --ont_r9_guppy5_sup --phased_output`

#### Variation and visualization statistics from VCF file

The vcftools and whatshap tools set
provide statistics on heterozygosity and visualizations for phased
haplotypes. Get stats for each and all contigs by
`whatshap stats PEPPER_MARGIN_DEEPVARIANT_FINAL_OUTPUT.phased.vcf.gz --tsv=pepper.phased.whatshap.tsv`

Visualizing in JBrowse.

1. Launch new session. Open sequence files. Name assembly “ant”. Select
   consensus fasta file and consensus fasta index (fasta.fai, created with
   the command `tabix file.fasta`).
2. Select “Launch View” and “Open”.
3. Select “Open Track Selector”. Click the “+” Button at bottom
   right.
4. Main file choose the variant file that aligned to the consensus
   (`PEPPER_MARGIN_DEEPVARIANT_FINAL_OUTPUT.haplotagged.bam`)
   and its index file (.bai). Confirm and add.
5. In track view, click the “three dot” menu next to the name of the
   bam file.
6. Select: Pileup settings -> sort by -> sort by tag. Type
   “HP”
7. Select: Pileup settings -> color scheme -> color by tag. Type
   “HP”

Transition Transversion ratio can be determined with the following
command  
`vcftools --TsTv-summary --gzvcf ../pepper-margin-deepvariant-out/input/pepper-margin-deepvariant-out/PEPPER_MARGIN_DEEPVARIANT_FINAL_OUTPUT.phased.vcf.gz`

#### BCFtools to make variant diploid fasta file

Until now, we have worked with only a haploid version of the genome.
We must generate a separate fasta file based on the consensus that swaps
out all the alternative alleles identified in the variant call file
(.vcf). By sequencing a single diploid individual, the variants all
represent the opposite haplotype from the other parental allele. The
following command will swap the variants into the consensus file.

`cat consensus.fasta | bcftools consensus deepvariant-out/PEPPER_MARGIN_DEEPVARIANT_FINAL_OUTPUT.phased.vcf.gz > haplotype2.consensus.fasta`

### DNA Methylation Analysis

Use megalodon
v2.4.2. Call methylation and hydroxymethylation at all cytosine
contexts (CpG + CH) Download basecalling neural network models for guppy
from the ONT Rerio
repository. Use megalodon with the 5mC + 5hmC all context model
v001.

1. I use megalodon with pip install (don’t forget to run “conda
   deactivate” if using megalodon from pip install).
   `megalodon /ant1/path/to/fast5 --outputs basecalls mappings mod_mappings mods per_read_mods --reference consensus.clean.fasta --devices all --processes 23 --output-directory megalodon-results --guppy-params " --use_tcp" --guppy-server-path /opt/ont/guppy/bin/guppy_basecall_server --guppy-config res_dna_r941_min_modbases_5mC_5hmC_v001.cfg`
2. Post process megalodon output Get *only* CG methylation sites
   `megalodon_extras modified_bases create_motif_bed --motif CG 0 --out-filename CG-motif.consensus.bed ../clean-assembly/consensus.clean.fasta`
3. Post process megalodon output Get non-CG (CH) methylation sites
   `megalodon_extras modified_bases create_motif_bed --motif CH 0 --out-filename CH-motif.consensus.bed ../clean-assembly/consensus.clean.fasta`
4. Intersect bed files to add 5mC values to CpG sites
   `bedtools intersect -a modified_bases.5mC.bed -b CG-motif.consensus.bed > CG-motif.5mC.methvalues.bed`
   `bedtools intersect -a modified_bases.5hmC.bed -b CG-motif.consensus.bed > CG-motif.5hmC.methvalues.bed`
5. Intersect bed files to add 5mC values to CH sites.
   `bedtools intersect -a modified_bases.5mC.bed -b CH-motif.consensus.bed > CH-motif.5mC.methvalues.bed`
6. Intersect bed files to add 5hmC to CpG sites
   `bedtools intersect -a modified_bases.5hmC.bed -b CG-motif.consensus.bed > CG-motif.5hmC.methvalues.bed`
   (last two columns are “depth of coverage” & “percentage of
   methylation”)
7. Repeat for CH sites.
8. Use awk to average the methylation column for whole genome
   methylation. This case uses only positions where the count is > 10
   reads.
   `awk '$10>10 {total+=$11; count++} END{print total/count}' CG-motif.5mC.methvalues.bed`

### Commensal assemblies

#### Wolbachia assembly

First align to a known wolbachia species such Wolbachia pipientis,
then create consensus with flye. Second, remap reads to flye-consensus,
and re-assembly with flye. Third, polish consensus.

1. Map to known wolbachia species.  
   `minimap2 -ax map-ont -t 24 genomes/wolbachia_pipentis_ref/GCF_014107475.1_ASM1410747v1_genomic.fna ../ant-total.fq.gz > ant-reads.vs.wolbacha.pipientis.sam`
2. Filter for aligned reads and convert to fastq for flye
   input.  
   `samtools view -b -F 4 ant-reads.vs.wolbacha.pipientis.sam > ant-reads.vs.wolbacha.pipientis.aln.bam`
   `bamToFastq -i ant-reads.vs.wolbacha.pipentis.aln.bam -fq ant-reads.vs.wolbacha.pipentis.aln.fq`
3. Convert to fasta.
   `seqkit fq2fa <reads.fq> -o reads.fa`
4. Remove all empty entries.
   `awk 'BEGIN {RS = ">" ; FS = "\n" ; ORS = ""} {if ($2) print ">"$0}' input.fas > output.fas`.
5. Now you have to remove the duplicate entries.
   `awk '/^>/{f=!d[$1];d[$1]=1}f' in.fa > out.fa`

Flye will run successfully now.

5. Generate consensus with flye.
   `flye --nano-hq ant-reads.vs.wolbacha.pipientis.aln.full.nodups.fa --out-dir flye-wolbachia --genome-size 1185k --threads 12`
6. Polish consensus with Racon and Medaka as above.

#### Mitogenome assembly

First align to a related mitogenome, the fire ant mitogenome. Then
create consensus with flye. Second, remap reads to flye-consensus, and
re-assembly with flye.

1. Map to related mitogenome species.  
   `minimap2 -ax map-ont -t 24 genomes/fireant-mtDNA/Solenoposis_invicta_mtdna.fasta ../ant-total.fq.gz > ant-reads.vs.fireantmtDNA.sam`
2. Filter for aligned reads and convert to fastq for flye
   input.  
   `samtools view -b -F 4 ant-reads.vs.ant-reads.vs.fireantmtDNA.sam > ant-reads.vs.fireantmtDNA.aln.bam`
   `bamToFastq -i ant-reads.vs.fireantmtDNA.aln.bam -fq ant-reads.vs.fireantmtDNA.aln.fq`
3. Convert to fasta.
   `seqkit fq2fa <reads.fq> -o reads.fa`
4. Remove all empty entries.
   `awk 'BEGIN {RS = ">" ; FS = "\n" ; ORS = ""} {if ($2) print ">"$0}' input.fas > output.fas`
5. Now you have to remove the duplicate entries.
   `awk '/^>/{f=!d[$1];d[$1]=1}f' in.fa > out.fa`

   Flye will run successfully now.
6. Generate consensus with flye. Flye does not deal well with such
   extremely high coverage. Use the `--meta` flag to get an
   assembly for mitogenomes. Then it is able to assemble.
   `flye --nano-hq ant-reads.vs.ant-reads.vs.fireantmtDNA.aln.full.nodups.fa --out-dir flye-mitogenome --genome-size 16k --threads 12 --meta`
   This generated 12 contigs. Blast was used to determine that contig\_1 was
   the correct sequence (96% query cover & 91% identity to C. atrox
   mitochondria). Save output as
   `mitogenome.flye.consensus_16k.fasta`
7. Polish consensus. First realign the reads to the mitogenome
   consensus.
   `minimap2 -ax map-ont -t 14 mitogenome.flye.consensus_16k.fasta ../ant-total.fq.gz > ant-reads.vs.mitogenome.flye.consensus_16k.sam`.
   Then polish 2X with Racon (realigning in between).
   `racon --cudapoa-batches 40 -t 16 ../ant-total.fq.gz ant-reads.vs.mitogenome.flye.consensus_16k.sam mitogenome.flye.consensus_16k.fasta > mitogenome.flye.consensus_16k.racon1.fasta`

##### Mitogenome annotation and visualization.

Use the consensus mitogenome as input to MITOS2 for gene
identification. `mitogenome.flye.medaka.consensus.fasta`.
Re-order the fasta file by cut and paste to put the cox1 gene in
position 1 as is customary. Resubmit to MITOS2 for proper gene
coordinates. Use MITOS2 output .tbl file which is already in 5 column
NCBI format + the consensus fasta as in put to Galaxy at tamu.edu’s
tool, “Genome polishing and submission” -> Five column tabular to
Genbank” to create a .gb file. Remember to force the format for the .tbl
file to “tabular” since Galaxy doesn’t auto-detect it correctly. View
the .gb file with Open Vector Editor. Edit the .tbl or .gb file to fix
any annotation oddities.

Note: Had to manually remove a frameshift inserted ‘t’ at position
2840. Homopolymeric stretch of ‘t’.

#### Blochmannia assembly

Flye does not deal well with such extremely high coverage. For
Blochmannia use the `--asm-coverage 50` flag to use the
longest 50 reads. Then it was able to assemble.  
`flye --nano-hq ant-reads.vs.blochmannia.aln.full.dedup.fa --out-dir flye-blochmannia --genome-size 791k --threads 12 --asm-coverage 50`.
Then polish 2X with Racon (realigning in between) .

### Miscellaneous code for filtering sam files.

To convert directly from sam to sorted bam without writing
intermediates to disk:
`samtools view -u file.sam |samtools sort -@ 24 -o <file>.sorted.bam`

To filter for UNmapped reads
`samtools view -f 4 file.bam > unmapped.sam`

To filter for Mapped reads
`samtools view -b -F 4 file.bam > mapped.bam`

### Session info


```
sessionInfo()
```
